# Low base-substitution mutation rate in the germline genome of the ciliate *Tetrahymena thermophila*

**DOI:** 10.1101/025536

**Authors:** Hongan Long, David J. Winter, Allan Y.-C Chang, Way Sung, Steven H. Wu, Mariel Balboa, Ricardo B. R. Azevedo, Reed A. Cartwright, Michael Lynch, Rebecca A. Zufall

## Abstract

Mutation is the ultimate source of all genetic variation and is, therefore, central to evolutionary change. Previous work on Paramecium tetraurelia found an unusually low germline base-substitution mutation rate in this ciliate. Here, we tested the generality of this result among ciliates using Tetrahymena thermophila. We sequenced the genomes of 10 lines of T. thermophila that had each undergone approximately 1,000 generations of mutation accumulation (MA). We applied an existing mutation-calling pipeline and developed a new probabilistic mutation detection approach that directly models the design of an MA experiment and accommodates the noise introduced by mismapped reads. Our probabilistic mutation-calling method provides a straightforward way of estimating the number of sites at which a mutation could have been called if one was present, providing the denominator for our mutation rate calculations. From these methods, we find that *T. thermophila* has a germline base-substitution mutation rate of 7.61 × 10^−12^ per site, per cell division, which is consistent with the low base-substitution mutation rate in P. tetraurelia. Over the course of the evolution experiment, genomic exclusion lines derived from the MA lines experienced a fitness decline that cannot be accounted for by germline base-substitution mutations alone, suggesting that other genetic or epigenetic factors must be involved. Because selection can only operate to reduce mutation rates based upon the “visible” mutational load, asexual reproduction with a transcriptionally silent germline may allow ciliates to evolve extremely low germline mutation rates.

## INTRODUCTION

Mutation is the ultimate source of all genetic variation, and the rate, molecular spectrum, and phenotypic consequences of new mutations are all important drivers of biological processes such as adaptation, the evolution of sex, the maintenance of genetic variation, aging, and cancer. However, because mutations are rare, detecting them is difficult, often requiring the comparison of genotypes that have diverged from a common ancestor by at least hundreds or thousands of generations. Further, interpreting the results of such comparisons is complicated by the fact that mutations are frequently eliminated by natural selection before they can be studied.

Mutation accumulation (MA) is a standard method for studying mutations experimentally. In a typical MA experiment, many inbred or clonal lines are isolated and passed repeatedly through bottlenecks. This reduces the effective population size and lessens the efficiency of selection, allowing all but the most deleterious mutations to drift to fixation (Bateman 1959; Mukai 1964). The genome-wide mutation rate and mutational spectrum can then be estimated by comparing the genomes of MA lines with those of their ancestors. Such direct estimates of mutational parameters are now available for a number of model organisms (Denver et al. 2009; Keightley 2009; Keightley et al. 2014; Lee et al. 2012; Lind and Andersson 2008; Lynch et al. 2008; Ness et al. 2012; Ossowski et al. 2010; Sung et al. 2012b; Zhu et al. 2014). However, the narrow phylogenetic sampling of these species still limits our ability to understand how mutation rates and patterns have evolved and, in turn, have influenced evolution across the tree of life.

Microbial eukaryotes are an extraordinarily diverse group, containing many evolutionarily distant lineages, some of which have unusual life histories and genome features (Katz and Bhattacharya 2006). However, microbial eukaryotes are often unsuitable for use in mutational studies because they are difficult to culture in the lab, especially at the small population sizes required to reduce the efficiency of selection in MA experiments. In addition, genomic resources (e.g., completed annotated reference genomes) are lacking for most eukaryotic microbes. These barriers have limited MA experiments to well-annotated model microbial eukaryotes such as *Saccharomyces cerevisiae* (Lynch et al. 2008; Zhu et al. 2014), *Schizosaccharomyces pombe* (Behringer and Hall 2015; Farlow et al. 2015), *Paramecium tetraurelia* (Sung et al. 2012b), *Dictyostelium discoideum* (Saxer et al. 2012), and *Chlamydomonas reinhardtii* (Ness et al. 2012; Ness et al. 2015; Sung et al. 2012a; fig. 1).

**Fig 1.**
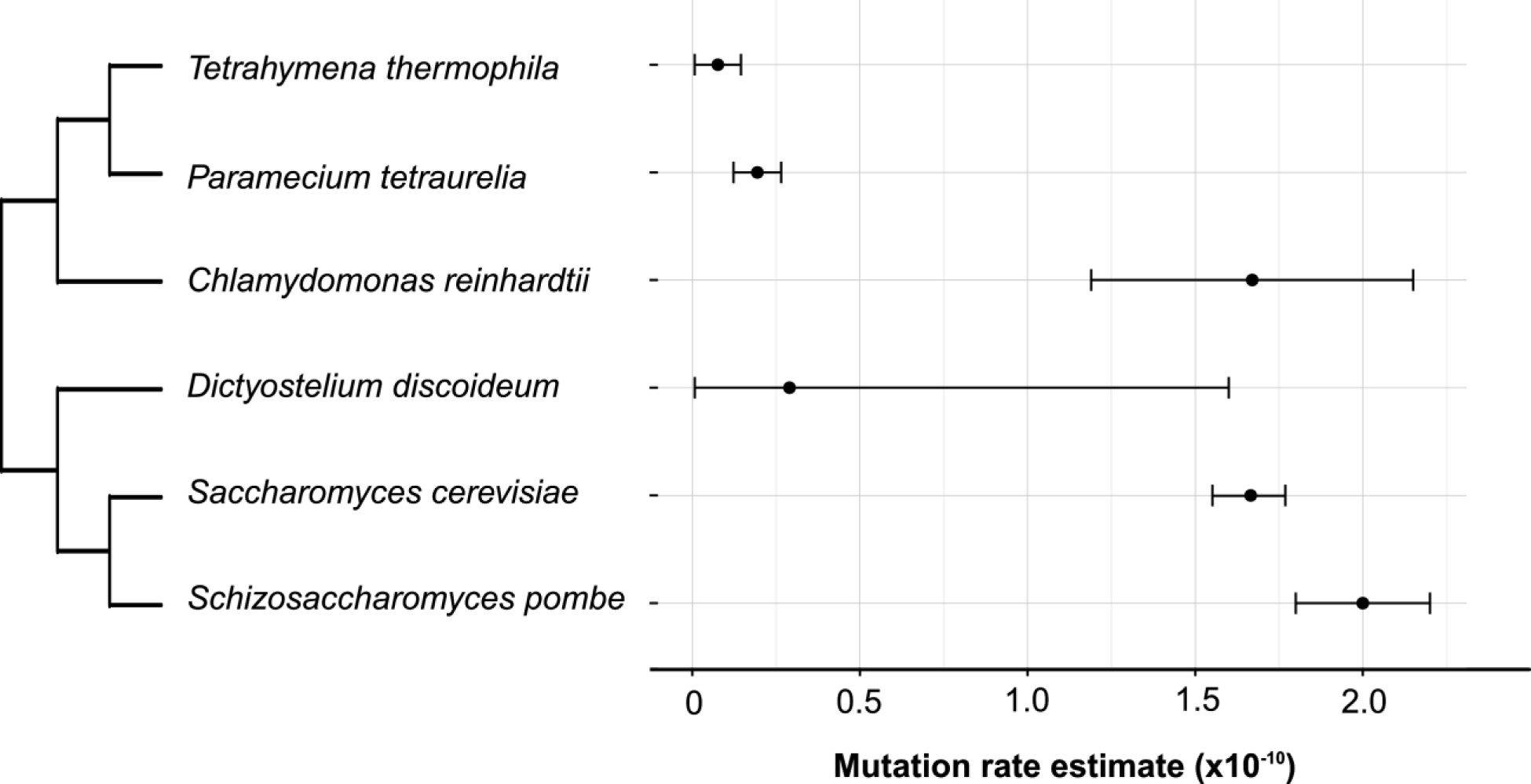
—Mutation rate estimates for unicellular eukaryotes. Base-substitution mutation rates per nucleotide per generation estimated for different unicellular eukaryotes: *T. thermophila* (this paper), *P. tetraurelia* (Sung et al. 2012b), *C. reinhardtii* (Ness et al. 2015), *D. discoideum* (Saxer et al. 2012), *Sa. cerevisiae* (Zhu et al. 2014), and *Sc. pombe* (Farlow et al. 2015). The phylogenetic tree was retrieved from the Open Tree of Life (Hinchliff et al. 2015); branch lengths are arbitrary. Error bars are 95% confidence intervals calculated by ignoring uncertainty in the number of sites at which a mutation could be detected and the number of generations that each MA ran experiment ran for.

The ciliated unicellular eukaryote *Tetrahymena thermophila* is well suited to MA experiments. Like all ciliates, individuals from this species have distinct germline and somatic copies of their nuclear genome. During asexual growth, the contents of the germline genome are duplicated mitotically but are neither expressed nor used to generate a new somatic genome. But unlike most other ciliates (including *P. tetraurelia*, which senesces in the absence of periodic mating or autogamy), *T. thermophila* can be propagated this way indefinitely. Thus, during periods of asexual growth — which can last over thousands of generations (Doerder 2014) — mutations can accumulate in the germline genome without being exposed to natural selection, which is operating on the somatic genome. Long et al. (2013) confirmed that MA lines of *T*. *thermophila* can be propagated asexually for at least 1,000 generations and inferred that they accumulate mutations in their germline genomes with detectable effects on fitness after the mutations are expressed in the somatic genome. However, Long et al. (2013) did not estimate the mutation rate directly at the molecular level.

The only other existing MA experiment from a ciliate was performed on *Paramecium tetraurelia* (Sung et al. 2012b) and yielded the lowest known base-substitution mutation rate in a eukaryote. Sung et al. (2012b) suggested that this exceptionally low mutation rate is the result of the unusual life history of ciliates, in which a transcriptionally silent germline genome undergoes multiple rounds of cell division between sexual cycles. Measurement of the mutation rate of *T. thermophila* will help reveal whether a low mutation rate is a general feature of ciliates. In addition, natural populations of *T. thermophila* have been the focus of population-genetic studies (Katz et al. 2006; Zufall et al. 2013), so mutational parameters estimated from MA experiments can be related to population and evolutionary processes.

Although the life history of *T. thermophila* is ideal for MA experiments, some features of its genome complicate typical computational approaches to detecting mutations from short-read sequencing. The genome is extremely AT-rich (~78% AT), contains many low complexity and repetitive elements, and has an incomplete reference genome (Eisen et al. 2006). These features make mapping sequencing reads to the reference genome difficult, which may lead to false positives when using naive mutation detection methods. To overcome these difficulties, we have developed a novel probabilistic mutation detection approach that directly models the design of an MA experiment and accommodates the noise introduced by mismapped reads. We used both our new method and an existing mutation-calling pipeline (Sung et al. 2012b) to analyse the MA sequences.

Here we expand the work presented by Long et al. (2013) by directly estimating the base-substitution mutation rate in *T. thermophila*. Our results are consistent with the exceptionally low rate estimated for *P. tetraurelia*, indicating that low germline mutation rates may be a general feature of ciliates. We also use our estimated rate to calculate the effective population size of *T. thermophila* in the wild. Our results establish that it is possible to estimate the mutation rate of *T. thermophila* directly from sequence data, but owing to the extraordinarily low rate, longer and larger MA experiments will be required to confidently estimate the mutational spectrum of a species with such a low mutation rate.

## MATERIALS AND METHODS

### Cell lines

The 10 evolved cell lines that were used in this study were generated from 10 parental MA lines (Supplementary Table S1). These lines were established from a single cell of the strain SB210 as described in Long et al. (2013). Briefly, the 10 MA lines were cultured in rich medium (SSP) in test tubes (Gorovsky et al. 1975) and experienced ~50 single-cell bottlenecks and ~1000 cell divisions, except for M28, which was bottlenecked 10 times and passed ~200 cell divisions. The optical density of cultures was measured prior to each transfer and the number of generations calculated using a standard curve of optical density for the ancestor (Long et al. 2013). Because directly sequencing the *T. thermophila* micronuclear genome is not feasible, we generated autozygous lines with macronuclear genomes derived from haploid copies of our ancestral and descendant micronuclear genomes using genomic exclusion (Allen 1963). Genomic exclusion lines were produced by two rounds of crossing between the MA lines (mating type VI) and a germline-dysfunctional B* strain (mating type VII, Bruns and Cassidy-Hanley, 1999). A mutation in the macronuclear genome of a genomic exclusion line derived from an MA line is assumed to correspond to a germline mutation in that MA line.

We accounted for heterozygosity in the ancestral strain by generating 19 independent genomic exclusion lines from the progenitor line. The DNA from all 19 genomic exclusion lines was pooled before library construction, allowing us to sequence both alleles at any heterozygous sites.

### Whole-genome sequencing

DNA libraries with insert size ~350 bp were constructed and Illumina paired-end sequenced by the DNASU core facility at the Biodesign Institute at Arizona State University and the Hubbard Center for Genome Studies, University of New Hampshire. The mean sequencing depth is ~47×, with >90% of the sites in the genome covered in all the sequenced lines (Supplementary Table S1). Sequencing reads are available from the NCBI’s SRA database under a BioProject with accession number PRJNA285268.

### Base-substitution analysis

We used two independent approaches to call point-mutations to avoid false negatives that might not be detected by a single approach. First, a widely used consensus approach (Sung et al. 2012b). Second, a probabilistic approach that adapts methods designed for family-based data to the design of MA experiments (Cartwright et al. 2012). Our list of candidates was generated by the union of calls from both methods.

#### Consensus approach

For the consensus approach we applied the following filters to reduce false positives that may arise from sequencing, read mismapping or library amplification errors. (1) Two mapping programs, BWA 0.7.10 (Li and Durbin 2009) and novoalign (V2.08.01; NOVOCRAFT Inc), were used in two independent pipelines to reduce algorithm-specific read mapping errors. (2) Only uniquely mapped reads were used (BWA option: sampe –n 1; NOVOCRAFT option: novoalign –r None), with mapping/sequencing quality scores > 20 (samtools mpileup –Q 20 –q 20). (3) The line-specific consensus nucleotide at a genomic site needed support from greater than 80% of reads to filter out false positives from mismapping of paralog reads. (4) Three forward and three reverse reads were required to determine the line-specific consensus nucleotide, to reduce false positive calls due to errors in library construction or sequencing. Putative mutations were then called if a single line was different from the consensus of all the remaining lines following the approach of Sung et al. (2012b). This approach has been applied to a wide variety of prokaryotic and eukaryotic organisms and repeatedly verified with Sanger sequencing (Denver et al. 2009; Lee et al. 2012; Long et al. 2015; Ossowski et al. 2010; Sung et al. 2015). The consensus approach also makes predictions consistent with those of the GATK SNP caller (Behringer and Hall 2015; Farlow et al. 2015).

#### Probabilistic approach using accuMUlate

The challenge of identifying mutations from genomic alignments can also be treated as a hidden-data problem (Cartwright et al. 2012). Fig. 2 illustrates the application of a hidden-data approach to our MA experiment. For a given site in the reference genome, the only data we observe directly is the set of sequencing reads mapped to that site. In order to determine if a mutation has occurred at the site, we have to consider the processes by which the read data was generated. These processes include biological processes (e.g., inheritance, mutation, genomic exclusion) and experimental processes that can introduce errors (e.g., library preparation, sequencing, data processing). Because none of these states are directly observed, we consider them to be hidden data. Each unique combination of hidden states represents a distinct history that could have generated the read data for a given state. See fig. 3 for an example of one such history with hidden and observed data illustrated.

**Fig 2.**
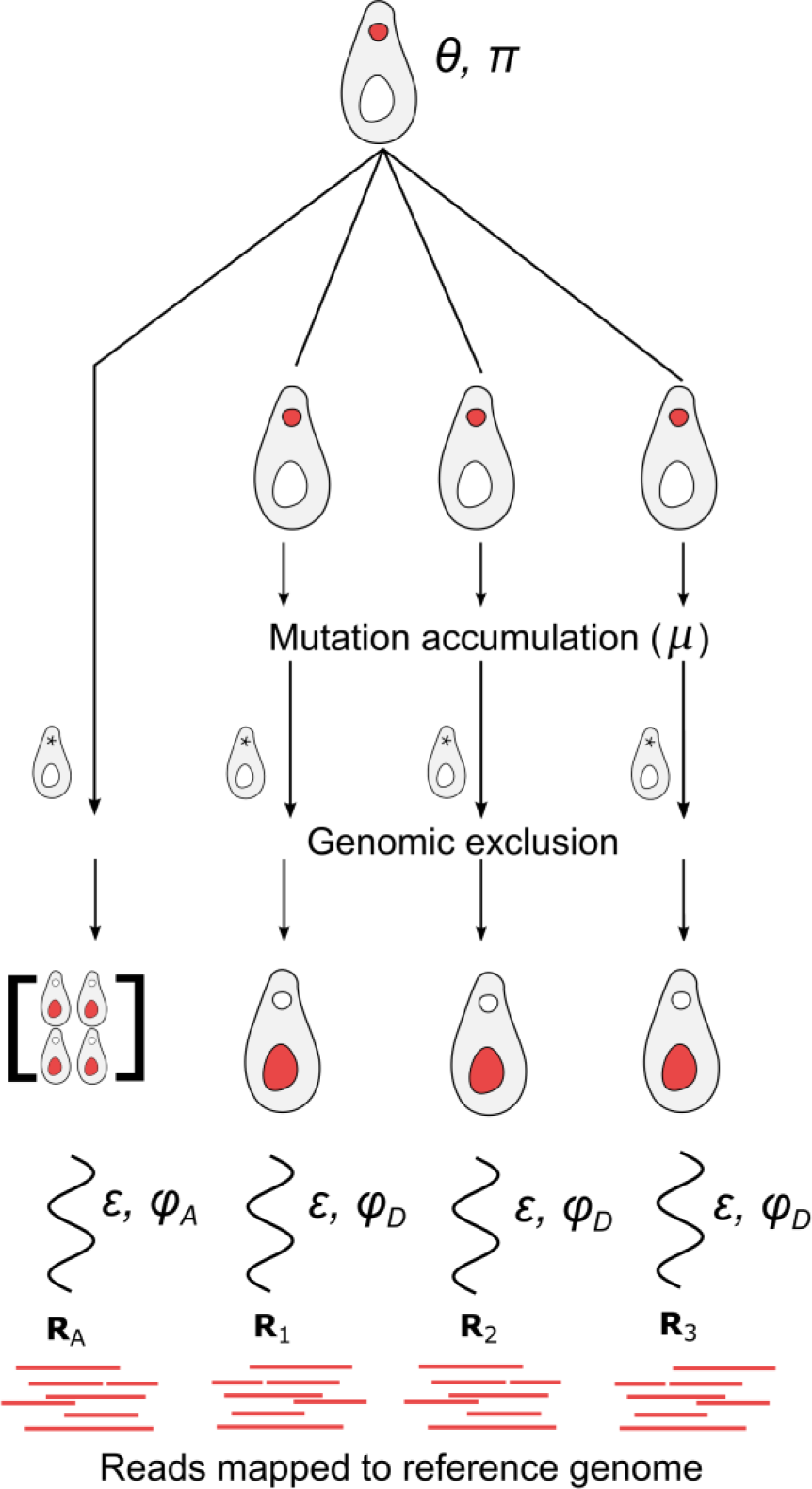
—Experimental design in relation to parameters of probabilistic mutation-detection model. A complete description of the experiment is presented in Long et al. (2013). Here, we describe how the experiment relates to the parameters used in our probabilistic mutation-calling model. Specifically, the ancestral line with average heterozygosity *θ* and genome-wide nucleotide frequencies *π* is used to generate a set of MA lines. Each of these lines accumulates mutations at a rate *μ* per nucleotide-site per generation for 1000 generations. Genomic exclusion, an auto-diploidization process, is used to generate lines with macronuclei representing one haploid-copy of each MA line (and multiple copies of the ancestral line, in order to detect ancestral heterozygosity). The macronuclear genomes of these genomic exclusion lines are then sequenced with a sequencing error rate of *ε* and overdispersion caused by library preparation and other correlated errors modeled as *φ*_*A*_ and *φ*_*D*_ for ancestral and descendant lines respectively. A full description of this model and its parameters is given in the subsection of the Materials and Methods labeled “Probabilistic approach using accuMUlate”.

**Fig 3.**
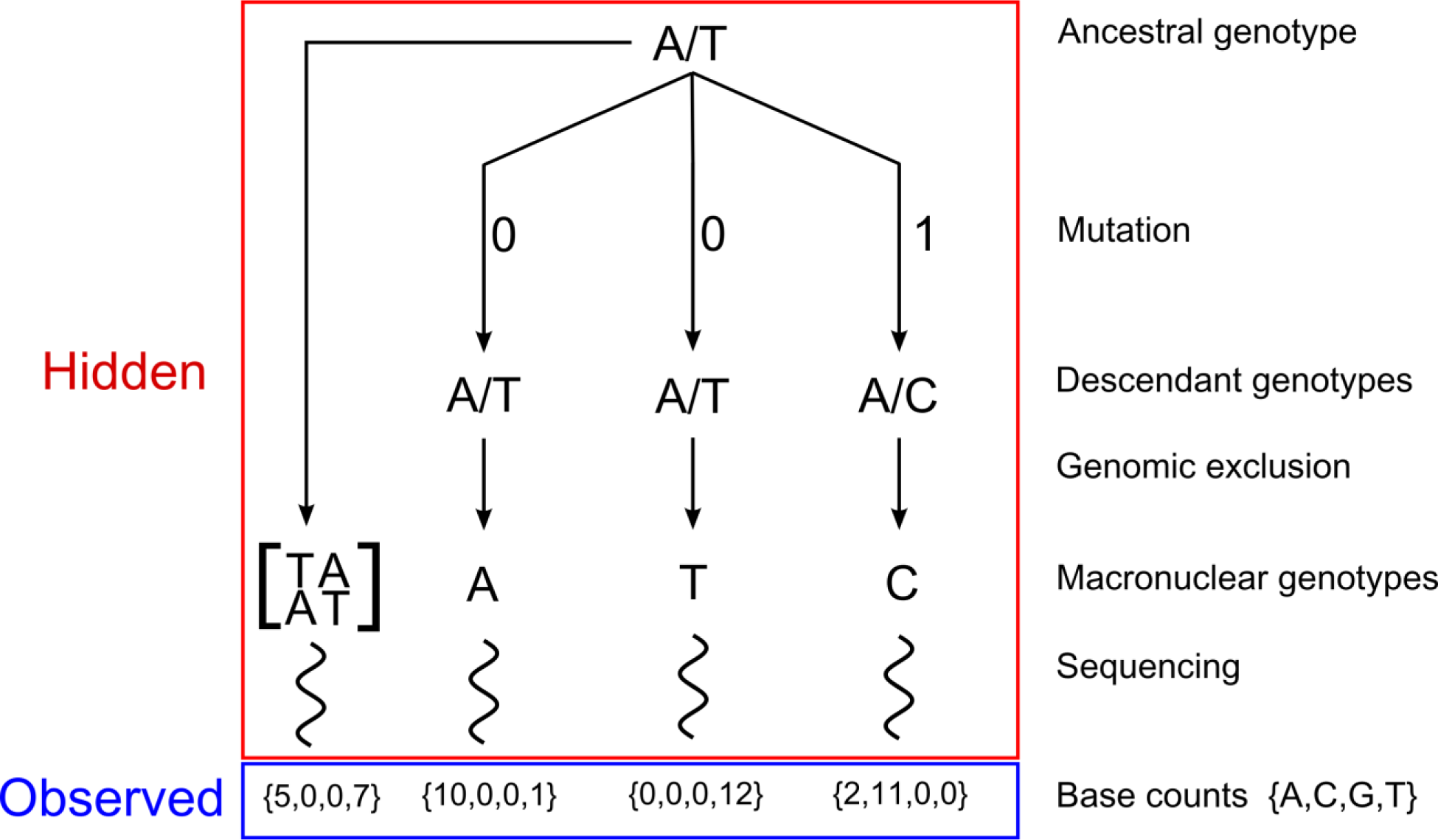
—Illustration of a single history in the accuMUlate method. In our model, a history is a unique combination of states (i.e. the genotypes of ancestral and MA lines, results of genomic exclusions and errors introduced during sequencing) generated during an MA experiment. Here we illustrate one such history by giving values to the different states in a model reflecting the same experimental design as fig. 2 and show how we calculate the probability that this history occurred and generated the observed sequencing data. Because we treat sites in the reference genome independently, we describe the process for a single site. Specifically, we consider a history in which an ancestor that is heterozygous with genotype A/T is used to establish three MA lines. One of those lines experiences a mutation from A/T to A/C, and the C allele of this mutant is passed on to a new macronuclear genome through genomic exclusion. The only data we observe for this locus is the set of bases mapped to this site that pass our filtering steps. We represent this data as vectors containing the number of A, C, G and T bases mapped to a given site (the base counts). We can use Equation 3 to calculate the probability that this sequencing data was generated by the specific history shown here. To do this, we first calculate the probability that the ancestor would have genotype A/T and that the observed sequencing data from the ancestor could be generated from this genotype (using Equations 6 and 4, respectively). Next, we consider the MA (descendant) lines, calculating the probability that the three descendant lines would have genotypes A, T and C and that the observed sequencing data could be generated from these genotypes. In this case we use the Felsenstein (1981) model of nucleotide substitution to calculate the probabilities that genomic exclusions generated from the MA lines would have these genotypes. We use the same genotype likelihood model (Equation 4) to calculate the probability that the sequencing data was generated from MA lines with these genotypes. Because each of the descendant lines is independent of each other, the overall probability of the history is simply the product of the probabilities for the ancestral and all descendant lines (Equation 3). We calculate the probability of a site containing at least one mutation by repeating this procedure for all possible histories at a given site (i.e. all possible combinations of genotypes) and keeping track of those histories that contain one or more mutations (Equation 2)

With the above formulation, our challenge is to determine the probability that a site contains at least one *de novo* mutation using our sequencing data (*R*) as the only observed input

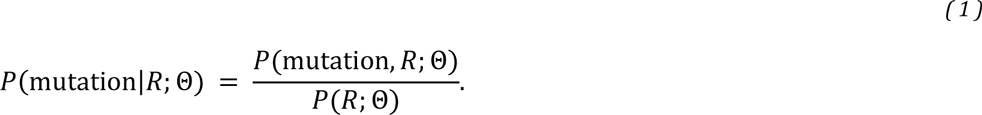

Here, *P*(mutation, *R*; Θ) is the joint marginal probability of at least one mutation being present and the sequencing data, and *P*(*R*; Θ) is the marginal probability of the sequencing data. The parameter Θ represents the model parameters and consists of the following:

- ***θ***, the proportion of sites in the ancestor that are heterozygous, approximately (see Equation 6);
- ***φ*_*A*_**, the overdispersion parameter for sequencing of the ancestor (described below);
- ***φ*_*D*_**, the overdispersion parameter for sequencing of the descendant lines (described below);
- ***π***, a vector representing the frequency of each nucleotide in the ancestral genome;
- ***μ***, the experiment-long mutation rate per site;
- ***ε***, the rate of sequencing error per site.

The numerator and denominator in Equation 1 are marginal probabilities. To calculate them from the full data we have to sum the probability of mutation over the full set of histories (*H*). Each of these histories is a unique combination of hidden states that could have generated the read data (an example of one such history is shown in fig. 3),

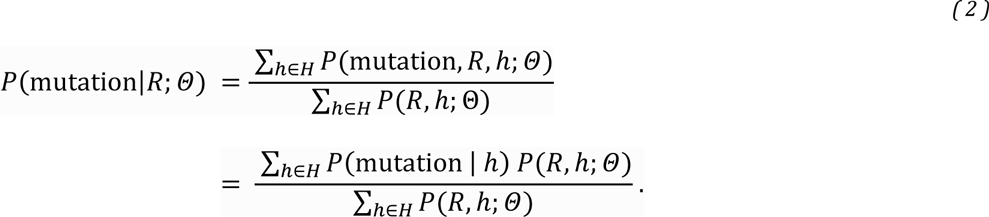

Note that the probability of there being at least one mutation in a given history *h*, *P*(mutation | *h*), is known to be either 1 or 0. Therefore, we only need to calculate *P*(*R*, *h*; Θ), the probability of the full data for the set of model parameters. This amounts to finding the probability that the read data was generated from an ancestral genotype *G_A_* that gave rise to descendants with genotypes specified by the particular history being considered. This can be calculated as the products of the prior probability of genotypes and the likelihoods of those genotypes given the set of all sequencing data (*R*),

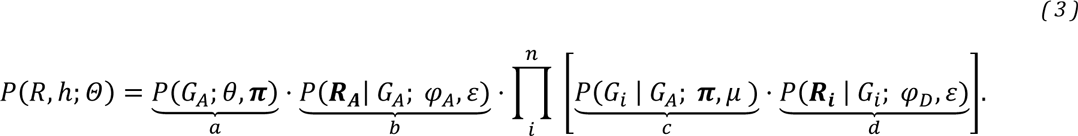

The elements of *R* are vectors of size four, and each contains the number of A, C, G and T bases mapped to this site for a particular sample (the pileup data). The specific element ***R_A_*** represents base counts from the ancestral strain. ***R_i_*** and *G_i_* are the base counts and genotype of the *i*-th descendant lines, respectively, and *n* is the total number of descendants.

The terms labeled “b” and “d” in Equation 3 are the probabilities of the observed sequencing data for a given genotype (i.e. genotype likelihoods). We calculate these genotype likelihoods using a Dirichlet-multinomial (DM) distribution. The DM is a compound distribution in which event-probabilities, ***p***, of a multinomial distribution is a Dirichlet-distributed random vector. Using a compound distribution provides flexibility to model the complex sources of error in sequencing data. To make this property of our model explicit, we use a parameterization of the DM distribution where ***p*** is a vector of length four containing the expected proportion of reads matching each allele and *φ* is an overdispersion parameter with values in the interval [0,1]. Using this parameterization, the DM distribution is equivalent to a simple multinomial distribution when *φ* = 0 and becomes increasingly overdispersed (i.e. the 200 variance increases) as *φ* tends to 1.

We demonstrate the calculation of genotype likelihoods using the term for the ancestral genotype in Equation 3 (“b” term) as an example. To calculate *P*(***R_A_***|*G*_*A*_; *φ*_*A*_, *ε*), we use the probability mass function of the DM distribution

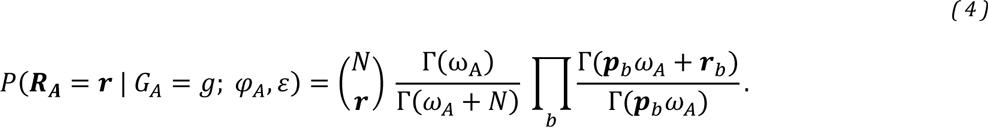

Here *N* is the total number of reads, 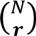 is the multinomial coefficient, Γ is the gamma function, and *ω*_*a*_ = (1 − *φ*_*A*_)/*φ*_*A*_. The parameter vector ***p*** contains the expected frequency of bases in {A, C, G, T} for the site under consideration and is indexed by *b*. Values in ***p*** are determined by both the probability of sequencing error (*ε*) and the ancestral diploid genotype *g* = {*g*_1_, *g*_2_} following Equation 5.

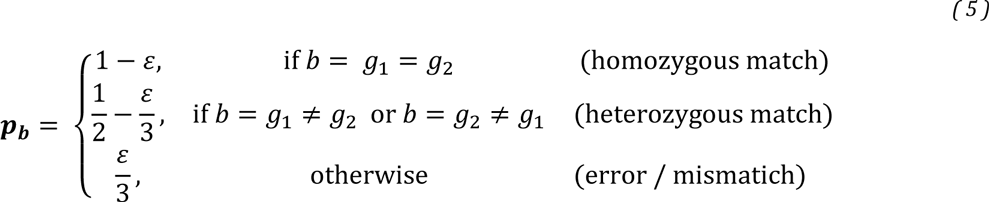

We now consider the remaining terms in Equation 3. The term labeled “a” represents the prior probability that the ancestor had a particular genotype (*g_A_* below) at the site under consideration given the nucleotide composition of the *T. thermophila* genome and average heterozygosity of the ancestral strain. We calculate this via a finite-sites model with parent-independent mutation,

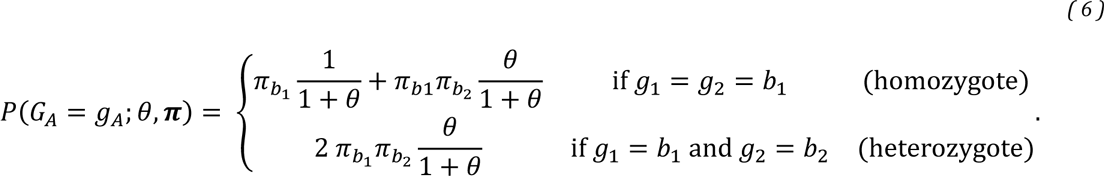

Here 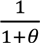 is the probability that the ancestor is autozygous at a site, and *π* is the vector of stationary nucleotide/allele frequencies in ancestral genome and *b*_1_ and *b*_2_ refer to the indices of the *g*_1_ and *g*_2_ alleles. See Wright (1949) for more details on this model and its biological assumptions.

To complete Equation 3 we need to consider the term labeled “c”, which represents the probability that the *i*-th MA line inherited a particular genotype, given the ancestral genotype and the probability of mutation. We calculate this via the Felsenstein (1981) model of nucleotide substitution. This model incorporates equilibrium nucleotide frequencies, allowing us to include the extreme AT-bias present in the *T. thermophila* genome.

Using the approach described above, we used Equation 2 to calculate both the probability of at least one point-mutation and the probability of exactly one point-mutation at every site along the *T. thermophila* reference genome. In MA experiments, multiple mutations at the same site are unlikely; therefore, sites that contain a strong signal of more than one mutation are likely false positives due to systematic errors in sequencing and mapping of reads.

This model is implemented in a C++ program called accuMUlate, which makes use of the Bamtools (Barnett et al. 2011) library. The source code used to perform the calculations described above is available under an MIT license from https://github.com/dwinter/accumulate; the specific version of the code used in these analyses is archived at http://dx.doi.org/10.5281/zenodo.19942. We ran our model on a genomic alignment produced by using Bowtie version 2.1.0 (Langmead and Salzberg 2012) to map reads to the December 2011 release of the *T. thermophila* macronuclear genome from the *Tetrahymena* Genome Database (Stover et al. 2006). One site in the reference contained a gap character, which we removed since our reads indicated that it was an artifact. We processed the resulting alignments to remove sequencing and mapping artifacts that could lead to false-positive mutation calls. In particular, we identified and marked duplicate reads using the MarkDuplicates tool from Picard 1.106 (http://pricard.sourceforge.net) and performed local realignment around potential indels using GATK 3.2 (DePristo et al. 2011; McKenna et al. 2010). We adjusted raw base quality scores by running GATK’s BaseRecalibrator tool, using a set of putative single nucleotide variants detected with SAMtools mpileup as input (Li et al. 2009).

The putative mutations from this approach were preliminarily identified by running accuMUlate to identify sites with a mutation probability > 0.1 with parameter-values: *φ*_*A*_ = *φ*_*D*_ = 0.001, *ε* = 0.01, *μ* = 10^−8^, *θ* = 0.0001 and only considering reads with Phred-scaled mapping and base quality scores ≥ 13 (corresponding to an estimated 5% probability of error). The validation phase showed that false-positive mutations were frequently associated with poorly-mapped reads, low coverage regions surrounding deletions with respect to the reference genome, or the presence of rare bases in all samples. Thus, we re-ran the accuMUlate model, excluding all reads with a mapping quality < 25 (corresponding to an estimated 0.3% chance of error), and using the overdispersion parameters *φ*_*A*_ = 0.03 and *φ*_*D*_ = 0.01. Setting *φ*_*A*_ > *φ*_*D*_ allowed us to accommodate the increased variance generated by sequencing pooled genomic exclusion lines to infer the ancestral genotype. In addition, we filtered out putative mutations that were not supported by at least 3 reads in both forward and reverse orientation. This final filtering step removed sites with unusually low coverage and those displaying strand bias, both characteristics associated with mismapped reads. We investigated the influence of our over-dispersion parameters by calculating the overall likelihood of the data using the initial and final parameter values. In order to make these results directly comparable, these calculations were performed on a data set consisting of all bases with quality score ≥ 13 from reads with a mapping quality ≥ 25.

### Validation of putative mutations

The validity of a subset of putative mutations was tested by Sanger sequencing. All mutations identified by either the consensus or the probabilistic approach were tested with suitable primers up to 500 bp away from the mutation site. Primers were designed using the default parameters of Primer3 (Koressaar and Remm 2007; Untergrasser et al. 2012) as implemented in Geneious (Kearse et al. 2012). Successful PCR products were purified and directly sequenced at Lone Star Labs (Houston, TX).

### Mutation rate calculations

Our probabilistic approach to mutation detection also provides a way to calculate the number of sites at which we could have detected a mutation if one was present, and thus the correct denominator to use for mutation rate calculations. Using our final model parameters, we shuffled the vector of read-counts generated from a given sample in order to simulate mutations in our data. This procedure was repeated for every site in the reference genome, shuffling the read counts from each descendent separately then recalculating the probability of a mutation. A site was treated as missing from a sample if the mutation probability calculated from shuffled read-counts was < 0.1 or if the most probable mutant allele was not supported by at least 3 reads in both the forward and the reverse orientation. To investigate the impact of our final parameter values and filtering criteria on the number of callable sites we repeated this procedure using the initial parameter set (i.e. with *φ*_*A*_ = *φ*_*D*_ = 0.001 and removing reads with mapping quality < 13). The number of callable sites detected using this approach for each line is given in Table 1.

**Table 1:**
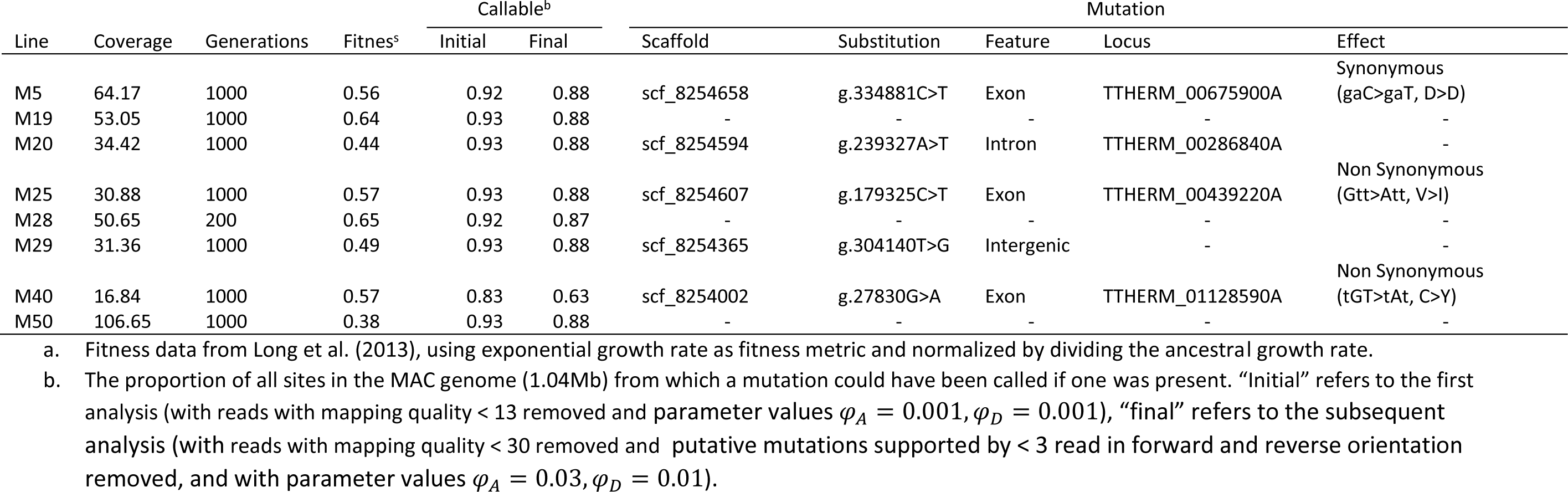
Summary of sequencing data and detected mutations. Note, no mutations were detected from lines M50, M28 or M19.

We calculated the mutation rate by summing the number of validated mutations (*n*_*i*_) across MA lines, and then dividing it by the product of the number of analyzed sites (*L*) and the number of generations (*T*) in each MA line (*i*): *û* = ∑_*n*_*n_i_*/(*LT*). Assuming that the number of mutations in each line follows a Poisson distribution (but not necessarily the same distribution) and ignoring uncertainty in our estimates for L and T, the standard error for our estimate of mutation rate was estimated as 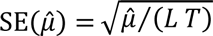, and a 95% confidence interval was constructed as *û*, ± 1.96 *SE*(*û*). We also performed the same calculation on log-transformed values of *μ*, *L*, and, *T* to produce a “log-space” confidence interval. To calculate genomic mutation rates we assumed a haploid genome size of 104 Mb (Eisen et al. 2006).

### Annotation of mutations

We annotated the functional context of identified mutations using snpEff (Cingolani et al. 2012) and the December 2011 release of the *T. thermophila* macronuclear genome annotation file from the *Tetrahymena* Genome Database.

## RESULTS

### Mutation detection and validation

To estimate the micronuclear mutation rate, we sequenced the whole macronuclear genomes of 10 homozygous genomic-exclusion lines, each derived from a separate *T. thermophila* line that had undergone MA for approximately 1000 generations. Using two different mutation-detection approaches (a widely used “consensus” method and a new probabilistic approach described in the Materials and Methods), we identified 92 sites for which there was some evidence of a mutation in at least one lineage. On closer inspection we found an unusual pattern — more than half of the apparent mutations were from lines M47 and M51, and in many cases reads containing the apparent mutant allele from one of these lines were also sequenced from the other line (but absent or very rare in all other lines).

To investigate this anomaly further we analyzed the frequency of non-reference bases in all samples across the whole genome (Supplementary Data). These analyses demonstrated that M47 and M51 differ from all other lines in the frequency of non-reference bases and in patterns of sequencing coverage. We do not know what caused the anomaly. It is possible that some cellular process occurred in these lines but not others (e.g., the incorporation of sequences usually restricted to the micronucleus, or the inclusion of DNA from the B* strain during genomic exclusion). It is extremely unlikely that M47 and M51 independently accrued more shared mutations than independent mutations during our MA experiment. For this reason, we have excluded these lines from all subsequent analyses.

Forty putative mutations remained after lines M47 and M51 were removed (Supplementary Table S2). We attempted to validate each of these mutations using Sanger sequencing. Only 4 of these mutations were validated. The remaining sites were either shown to be false positives (11 sites) or failed to generate either PCR amplicons or clean sequence traces (25 sites). Closer inspection of the data underpinning both the false positive and inconclusive mutations showed these sites to have unusually low sequencing coverage and low mapping quality, and to be subject to strand bias. All of these properties are associated with mapping error, and are known to generate false positive variant calls (Li 2014). For this reason, we re-ran our probabilistic mutation caller using stricter filters for mapping quality and excluding putative mutations that did not have at least 3 sequencing reads supporting a mutation in both the forward and reverse orientation. None of the inconclusive or false positive sites were called as mutations in this analysis, which also detected an additional mutation that was confirmed by Sanger sequencing. Thus, we detected a total of 5 mutations across 8 MA lines, with no line having more than one confirmed mutation (Table 1). Our probabilistic method produced more false positives than the consensus approach but generated no false negatives (Supplementary Table S2). Of the 5 mutations detected, 2 are non-synonymous, 2 are synonymous, and one is in an intergenic region.

### Number of callable sites

We estimated the denominator for our mutation rate estimates by calculating the number of sites at which a mutation could be called if one was present. An average of 86.1% of the reference genome was callable per line (Table 1). Sites for which we lacked power to detect mutations in at least one line are in relatively gene-poor regions; 30% of such sites are in exons compared to 49% of always-included sites. We also considered the impact of our final filtering steps and model parameters on our analyses. The more stringent filtering steps we used to generate our final mutation set reduced the proportion of callable sites per line, with the median proportion of callable sites declining from 93% to 88% (Table 1). The final overdispersion values used in our probabilistic mutation caller produced a better fit to our data than the initial values, with the overall log likelihood improving by 8 × 10^4^.

### Mutation rate

Given the number of callable sites, the 5 mutations that we detected yield a base-substitution mutation rate estimate of 7.61 × 10^−12^ per base pair per asexual generation (95% confidence interval, CI = [0.691 × 10^−12^, 14.53 × 10^−12^] using the standard method or [4.68 × 10^−12^, 12.38 × 10^−12^] using log-transformed values). This point estimate is approximately one third of the rate reported for *P. tetraurelia*, although the 95% CIs of both estimates overlap (fig. 1), and equates to a genome-wide rate of 0.8 base-substitution mutations per haploid genome per thousand asexual generations (95% CI = [0.07, 1.50]).

If our estimate of the base-substitution mutation rate holds for the portions of the genome from which we did not have sufficient power to detect mutations, then we estimate that we have failed to detect an additional 0.87 mutations across all of the macronuclear genomes sequenced.

## DISCUSSION

We have used whole-genome sequencing and a novel mutation-detection approach to estimate the base-substitution mutation rate of *T. thermophila* from 8 MA lines (Long et al. 2013) and obtained an estimate of 7.61 × 10^−12^ mutations per-site per-generation. This is the lowest estimate of base-substitution mutation rate from a eukaryote (for surveys of mutation-rate estimates see fig. 1 and Sung et al 2012b), and indeed lower than that observed in any prokaryote. However, it is not significantly different from the rate in either the social amoeba *Dictyostelium discoideum* (Saxer et al. 2012) or the ciliate *P. tetraurelia* (Sung et al. 2012b). The fact that the two lowest mutation rates have been recorded in ciliates supports the hypothesis that ciliates in general have low germline mutation rates (Sung et al. 2012b).

Direct estimates of the mutation rate from MA lines can only be as accurate as the methods used to detect mutations. Our estimate of a low mutation rate in *T. thermophila* could conceivably result from a high rate of false-negative results. However, we believe that this is unlikely. Our approach to mutation detection was designed to maximize the sensitivity of our analyses. We initially applied lenient filters to our data and attempted to validate all putative mutations detected by two separate methods. Most of the putative mutations suggested by this initial analysis could not be validated by Sanger sequencing. For this reason, we developed filters and model-parameters that improved the specificity of our mutation-calling method (producing negligible mutation probabilities for all of our unconfirmed mutations, while still supporting our confirmed mutations). It is possible that this increased stringency also led us to miss mutations present in our descendant lines. To account for the possibility of such false negatives in our mutation rate estimates, we simulated mutations in our data. This allowed us to identify sites at which we would not be able to detect a mutation in a given line even if one was present. Sites for which we could not call a simulated mutation were not included in the denominator of our mutation rate calculation. Thus, we are confident that our mutation rate estimate is accurate, at least for the regions of the genome from which we could call mutations.

Our mutation rate estimate allows us to estimate the effective population size of *T. thermophila.* If we assume that silent sites in protein-coding genes are effectively neutral and under drift-mutation equilibrium, the population-level heterozygosity at silent sites (π_s_) has expected value 4*N_e_*µ, where *N_e_* is the effective population size, and µ is mutation rate per site per generation. Using the estimate *N_e_* × µ = 8 × 10^-4^ reported by Katz et al. (2006), if we assume that mutation rates in the germline and somatic genomes are equal, our *N_e_* estimate for *T. thermophila* is 1.12 × 10^8^, which is almost identical to that of *P. tetraurelia* (*N_e_* = 1.24 × 10^8^; Sung et al. 2012b). These estimates may seem surprising given the observations that *P. tetraurelia* is cosmopolitan and regularly isolated from different continents (Catania et al. 2009), while *T. thermophila* has a distribution limited to the eastern United States (Zufall et al. 2013). However, the relationship between census population size and genetic diversity (and therefore estimated *N_e_*) is not a simple one (Leffler et al. 2012; Lewontin 1974). In very large populations stochastic processes, including demographic events that prevent populations from reaching mutation-drift equlibrium (Haigh and Maynard Smith 1972; Leffler et al. 2012) and the effects of selection on sites linked to neutral variants (Gillespie 2001; Lynch 2007; Neher et al. 2013), limit genetic diversity across the whole genome. Regardless, the large effective population size estimated here suggests that selection will have considerable power in the evolution of *T. thermophila*.

The unusual genome structure and life history of ciliates may explain their low mutation rates. Sung et al. (2012a) argued that mutation rates are minimized to the extent made possible by the power of natural selection — the “drift barrier” hypothesis. Selection operates to reduce the mutation rate based on the “visible” mutational load, and mutations that accumulate in the germline genome in ciliates during asexual generations are not expressed and exposed to selection until they are incorporated in a new somatic genome following sexual reproduction. Thus, the mutation rate per selective event is equal to the mutation rate per asexual generation multiplied by the number of asexual generations between rounds of sexual reproduction. The low mutation rates reported for ciliates may have evolved naturally as a consequence of the many asexual generations in between bouts of sexual reproduction, combined with large effective population sizes that promote strong selection for low mutation rates.

Unlike *P. tetraurelia*, *T. thermophila* does not undergo senescence in the absence of sex, and we lack a good estimate for the frequency of sexual reproduction in natural populations (Doerder et al. 1995). Therefore, we cannot put an upper bound on the number of asexual generations between conjugation events. However, we can estimate a lower bound because cells arising from sexual reproduction enter a period of immaturity lasting ~50–100 divisions (Lynn and Doerder 2012). We know that the germline genome divides at least this many times without opportunity for selection on any newly acquired mutations. Using the immaturity period as a proxy for the frequency of sex gives an estimate of the base-substitution mutation rate of ~0.1 mutations per haploid genome per conjugation event — much closer to that of other eukaryotes per round of DNA replication (Sung et al. 2012b).

Most mutations with effects on fitness are deleterious, so the accumulation of mutations in the absence of selection is expected to lead to a reduction in organismal fitness (Bateman 1959; Halligan and Keightley 2009; Mukai 1964; Muller 1928). The fitness of a genomic exclusion line derived from an MA line of *T. thermophila* should, in part, reflect the germline mutations in that MA line. If most germline mutations are base-substitutions, the low germline base-substitution rate would lead us to predict modest effects on the fitness of the genomic exclusion lines we studied. Surprisingly, some of these lines experienced substantial fitness losses relative to the ancestor (Long et al. 2013). For example, we did not detect any base-substitution mutations in the line with largest observed loss in fitness (M50, w=0.38) (Table 1). It is unlikely that the fitness losses observed in these MA lines can be explained by other undetected single-base substitutions, as our mutation calling method had power to detect mutations in an average of 86.1% of the genome (Table 1) and the excluded portion of the genome is relatively gene poor. Rather, it seems likely the fitness of these lines is determined in part by indels and other structural variants that we did not include in this study. Furthermore, non-Mendelian patterns of inheritance could obscure the relationship between mutations and fitness. For example, the fitness of an individual line may be influenced by epigenetic processes, such as cortical inheritance (Sonneborn 1963) or small RNA guided genome rearrangement (Mochizuki and Gorovsky 2004).

Our probabilistic mutation calling method can directly model the overdispersion produced by modern sequencing techniques. Our initial run of this method used very low values for the overdispersion parameters and produced many false positive mutation calls. We believe that these false positives arose from sites with data that did not fit the expectation of relatively clean data that these low overdispersion values represent. Using data form the validation phase, we were able to show that increasing the overdispersion parameter values (and being more stringent about which sequencing reads were included in our analysis) improved the accuracy of our method such that it produced fewer false positives while still correctly identifying all validated mutations. In addition to improving the accuracy of our mutation calling, increasing the values of the overdispersion parameters substantially increased the fit of our model to the sequencing data. Although the small number of true mutations in this experiment prevents us from performing a more complete analysis, this study demonstrates that modeling overdispersion in sequencing data can improve mutation calling methods.

In this study we have established that it is possible to detect mutations in *T. thermophila* MA lines through short-read sequencing, and thus to directly study the nature of mutation in this model organism. Although we were able to show that *T. thermophila* shares a low mutation rate with *P. tetraurelia* (the only other ciliate for which a mutation rate has been directly estimated), there is still much to learn about mutation in this species. For instance, the unusual genome structure of ciliates presents a novel test of the drift-barrier hypothesis of mutation rate evolution (Sung et al. 2012a). If the mutation rates of the germline and somatic nuclei can evolve independently then we would expect the somatic mutation rate to be higher (i.e. more similar to the mutation rates of other eukaryotes) because somatic mutations are exposed to selection after each cell division. Furthermore, the small number of mutations accumulated over this experiment has prevented us from analyzing the spectrum of mutations arising in *T. thermophila* and determining the influence of mutational biases on genome evolution. Similarly, the few mutations that we detect seem inadequate to explain the observed losses of fitness during MA. Future studies using more MA lines evolving over longer periods and detecting indels and other structural variants accrued during MA will be needed to fully understand the effects of mutation and selection in *T. thermophila*.

## ACKNOWLEDGEMENTS

We thank Kristen Dimond, Robert Coyne, Tom Doak, Kale Dai, Adam Orr, and Rachel Schwartz for technical help and two anonymous reviewers for helpful comments. This study is funded by NIH R01GM101352 (RAZ, RBRA, RAC) and Multidisciplinary University Research Initiative award W911NF-09-1-0444 (ML) from the US Army Research Office, NIH grant R01GM036827 (ML) and National Science Foundation Grant MCB-1050161 (ML).

